# Distinct denitrifying phenotypes of predominant bacteria mould nitrous oxide metabolism in upland agricultural soils

**DOI:** 10.1101/2022.03.08.481735

**Authors:** Qiaoyu Wu, Mengmeng Ji, Siyu Yu, Ji Li, Xiaogang Wu, Xiaotang Ju, Binbin Liu, Xiaojun Zhang

## Abstract

Denitrifying nitrous oxide (N_2_O) emissions in agroecosystems result from variations in microbial composition and soil properties. However, the microbial mechanisms of differential N_2_O emissions in agricultural soils are less understood. Microcosm experiments of two types of Chinese farmland soil were conducted with nitrate (250 mg/kg) and a combination of glucose (1000 mg/kg) and nitrate, and a case with no addition was used as the control. The results show that N_2_O accumulation in black soil (BF) was significantly higher than that in fluvo-aquic soil (FF) independent of carbon and nitrogen supply. The abundance of denitrifying genes was significantly higher in FF, but the ratios of genes responsible for N_2_O production (*narG, nir*S, and *nirK*) to the gene for N_2_O reduction (*nosZ*) did not significantly differ between the two soils. However, the soils showed obvious discrepancies in denitrifying bacterial communities. High accumulation of N_2_O was verified by the isolates of *Rhodanobacter*, which is predominant in BF due to its truncated denitrifying genes and lack of N_2_O reduction capacity. The dominance of complete denitrifiers such as *Castellaniella* in FF led to a rapid reduction in N_2_O and reduced N_2_O accumulation, as demonstrated when its corresponding isolate was inoculated into both studied soils. Therefore, the different phenotypes of N_2_O metabolism of the distinct denitrifiers maintained in the two soils caused their differing N_2_O accumulation. This knowledge could guide the regulation of the denitrifying bacterial community and the phenotypes of N_2_O metabolism in agricultural soils to reduce N_2_O emissions.

## Introduction

The nitrogen cycle and the associated microbes play an important role in the sustainability of ecosystems (Ishii et al., 2011). N_2_O has drawn much attention as an important ozone-depleting substance (Ravishankara et al., 2009) and has global warming potential approximately 265 times greater than that of CO_2_ (Dou et al., 2016). Agricultural soils are a major source (60%) of anthropogenic nitrous oxide (N_2_O) emissions globally (Lopez-Aizpun et al., 2020). Denitrification is a major source of N_2_O in agricultural soil and the main process responsible for N loss to the atmosphere (Butterbach-Bahl et al., 2013), although the nitrification process contributes a substantial proportion in a specific soil (Huang et al., 2014). Denitrification, the stepwise reduction of NO_3_^−^ to N_2_, involves four reduction steps. The enzymes catalysing four stepwise are encoded by the *narG* or *napA* (nitrate reductases), *nirK* and *nirS* (nitrite reductases), *norB* (nitric oxide reductase, responsible for N_2_O generation), and *nosZ* (N_2_O reductases) genes (Richardson et al., 2009). Microorganisms mitigate N_2_O emissions by acting as N_2_O sinks and counteracting those that act as N_2_O sources, which have higher catalytic activity for N_2_O production than for N_2_O reduction (Lycus et al., 2017).

Many studies have investigated the factors influencing N_2_O emissions from soil. Rates of N_2_O emission from agroecosystems are often explained by variations in soil properties and are strongly influenced by the abundance and diversity of communities of denitrifying microbes (Hu et al., 2015; Domeignoz-Horta et al., 2018). Some studies have focused on how environmental factors such as fertilization (Yan et al., 2003), irrigation (Owens et al., 2016), biochar (Harter et al., 2014), carbon substrate availability (Butterbach-Bahl and Dannenmann, 2011) and pH (Liu et al., 2010) affect denitrifying microorganisms, which further trigger the production of N_2_O from soil. It is a challenge to elucidate the mechanism of N_2_O emissions from agricultural soils in different climatic zones (Cheng et al., 2004), although the molecular investigation of microbial communities has expanded our knowledge of denitrifiers. To date, the correlation between denitrifying functional genes and N_2_O flux has been inconsistent for different soils. For example, Ullah et al. found potential denitrification activity to be significantly positively correlated with a higher abundance of denitrifying functional genes in black soil (BF) than in fluvo-aquic soil (FF) (Ullah et al., 2020). Yang et al. found the abundance of the *narG, nirS* and *nirK* genes to be correlated with annual N_2_O emissions in intensively managed calcareous FF (Yang et al., 2017). However, another study indicates that changes in denitrifier gene abundance are not the main factors influencing N_2_O emissions (Gao et al., 2016). Moreover, there is no correlation between N_2_O emissions and functional gene abundance in fumigated FF and lateritic red soil (Fang et al., 2018). Instead, some studies showed a linkage between the taxonomic composition of soil denitrifying bacteria in farmland soil and the denitrification rate or N_2_O production (Rich et al., 2003; Ji et al., 2021).

Microbial communities in different soils are structured over the long term by distal controls, which include both environmental factors and biotic interactions (Wallenstein et al., 2006; Hallin et al., 2009; Yin et al., 2015). BF and FF are among the most widely distributed farmland soils in China. BF is mainly distributed in Northeast China and is considered one of the most fertile soils in China due to its high levels of organic matter. The BF region plays a unique role in maintaining crop yield in China (Xu et al., 2010). The North China Plain (NCP) lies in the alluvial plain of the Yellow River (Ju et al., 2009) and is dominated by FF. This region is characterised by low levels of soil organic carbon, poor soil structure and high pH (Wu et al., 2018; Zhu et al., 2019).

It is noteworthy that BF and FF are now being irrigated with the application of a large amount of fertilizer N with the aim of obtaining higher yields, which has profoundly degraded soil quality and led to substantial total N_2_O emissions in the two regions (Cui et al., 2008; Ju et al., 2009; Xu et al., 2016). Compared to1990-2003 levels, the annual growth rate of N_2_O emissions from farmland in China decreased significantly from 2004 to 2014, but the rate in Heilongjiang Province, where BF is widely distributed, showed a significant increase (Shang et al., 2019). A field study found approximately 1.3% of N fertilizer inputs is lost as N_2_O from BF in Northeast China (Liang et al., 2004), which is much greater than the average of 0.6% found for N-fertilized upland soils in China (Xing, 1998). The levels of accumulated N_2_O from BF are higher than those from FF under different fertilization conditions in pot experiments (Zhang et al., 2021). Zhang et al. (Zhang et al., 2021) found that *nirK* and *nosZ* gene-containing denitrifiers in BF and FF are distinct and that *nosZ* gene abundance is the only biological factor that significantly determines N_2_O emissions in different soils through a structural equational model. However, this study did not pay attention to the effects of differences in the composition of denitrifying bacteria in the two soils. Does this difference relate to the difference in N_2_O emissions from BF and FF? It is urgently necessary to unravel the underlying mechanisms causing differences in N_2_O accumulation in these two types of soil to establish an effective means to mitigate the N_2_O emission resulting from agricultural activity.

We hypothesize that the varied community contains denitrifiers with different metabolic phenotypes, which shapes the distinct denitrification processes in the soils. To test this hypothesis, in this study, we performed comparative studies with microcosm experiments using BF and FF under three controlled incubation conditions with different amounts of nitrate and glucose. We used a robotized incubation system to continuously measure N_2_O and N_2_ fluxes. Furthermore, we explored the structure of the total bacterial and denitrifying bacterial communities by high-throughput 16S rRNA gene sequencing and quantified key functional genes (*narG, nirS*/*K*, and *nosZ* for denitrifiers). Moreover, we isolated the dominant denitrifying bacteria from both soils and verified their denitrifying properties. As a result, this study explored the microbial mechanisms for different accumulation of N_2_O in the two main types of soils used in Chinese agriculture.

## Materials and methods

### Soil used for experiments

Two types of soil samples were collected from farmland and used for incubation experiments in this study. Black soil (BF) was collected from a long-term experimental field in Lishu County(43^°^18′N, 124^°^14′E), Jilin Province, located in Northeast China. Fluvo-aquic soil (FF) was collected from a long-term experimental field at the Shangzhuang experimental station of China Agricultural University (40^°^08′N, 116^°^10′E), located on the North China Plain. The cropping system and physicochemical properties of the original soil are shown in (Table 1). Samples of each type of soil were taken from five different points at depths ranging from 0-20 cm and mixed as the final sample. All samples were placed in black plastic bags and stored at 4 °C before use. Soil samples were sieved (2 mm) to remove stones and coarse roots prior to incubation experiments.

**Table 1.** Physicochemical properties and potential denitrification rate (PDR) in BF (black soil) and FF (fluvo-aquic soil). The data are expressed as mean ± SD.

|  | **BF** | **FF** |
| --- | --- | --- |
| Latitude and longitude | 43°18'N,124°14'E | 40°18'N,116°10'E |
| Cropping system | maize | wheat, maize |
| Manure input (kg N hm<sup>-2</sup> year<sup>-1</sup>) | N: 234 kg N /ha<br>P<sub>2</sub>O<sub>5</sub>: 108 kg/ha<br>K<sub>2</sub>O:108 kg/ha | Winter wheat 300 kg N/ha<br>Summer corn 260 kg N/ha |
| Nitrogen management | compound fertilizer | Conventional fertilization |
| pH | 7.9±0.1 | 8.3±0.1 |
| Water content (%) | 18.90±0.04 | 10.17±0.06 |
| WHC(%) | 45.26±1.50 | 41.34±0.08 |
| DOC | 26.95±1.11 | 18.08±0.96 |
| PDR (nmol g<sup>-1</sup> h<sup>-1</sup>) | 26.07±0.46 | 38.69±1.62 |
| NO<sub>2</sub><sup>-</sup> (mg/Kg) | 0 | 0 |
| NO<sub>3</sub><sup>-</sup> (mg/Kg) | 13.67±0.12 | 26.11±0.47 |
| NH<sub>4</sub><sup>+</sup> (mg/Kg) | 4.94±5.73 | 0.26±0.29 |

### Experimental design and soil incubation

Before the experiment was conducted, soil samples were pre-incubated at 25 °C in the dark for 7 days to stabilize the microbial activity and avoid pulse effect on soil respiration. Soil was transferred to 120 ml serum vials (20 g dry weight per vial) containing buffer solution according to the following experimental design. Soil samples underwent CK, N250, N250+G treatments (without any addition, with initial nitrate content of 250 mg/kg, and with initial nitrate content of 250 mg/kg plus glucose content of 1000 mg/kg, respectively) while controlling the final soil moisture to 70% WHC (water holding capacity). The triplicate vials containing soil samples were sealed with airtight butyl-rubber septa and aluminum crimp caps. The headspace of the serum vials was alternately evacuated and refilled with high purity helium (99.999%) four times to create a completely anaerobic environment without O_2_ and N_2_. All vials were balanced to atmospheric pressure and incubated under anaerobic conditions at 25 °C for 7 days.

### Measurement of N_2_O and N_2_ fluxes and soil parameter

During 7 days’ anaerobic incubation, the headspace of serum vials was sampled every 4 hours and analyzed for N_2_ and N_2_O concentration in a robotized incubation system (ROBOT) similar to that described by Molstad (Molstad et al., 2007), with moderate improvements. The gas samples were drawn by a peristaltic pump, which returned an equal volume of helium. To characterize the ratio of N_2_O in a mixture of nitrogen elements, we calculated an N_2_O production index (*I*_N2O_) of N_2_O/(N_2_O+N_2_) during anaerobic incubations as described in previous literature (Liu et al., 2010). The index was calculated for each vial and used as single observation in the statistical analyses.

Soil pH was measured in a 1:5 (w/v) soil to H_2_O slurry mixture with a pH meter (Mettler-Toledo, Switzerland). DOC was monitored by a total organic carbon analyzer (TOC-LCSH, Shimadzu, Japan). Soil nitrate content was extracted using 1 M KCl solution at a 1:10 (w/v) soil to H_2_O and determined using an automatic chemical discontinuity analyzer. The content of ammonium in soil was estimated by indophenol blue method. Soil nitrite was determined by N-(1-naphthyl)-ethylenediamine dihydrochloride spectrophotometric method (Kesari and Gupta, 1998). The gravimetric soil water content was determined by the oven drying method at 105 °C for 24 h. The total of the N_2_O and N_2_ emission rate was calculated as potential denitrification rate (PDR) using the slope of the linear regression between the changes in concentration of either N_2_O or N_2_ versus time (Qin et al., 2017).

### Soil DNA extraction and quantification of denitrifying genes

Total DNA was extracted from 0.3 g of frozen soil sample according to CTAB (hexadecyl trimethyl ammonium bromide) method (Paulin et al., 2013). DNA from soil samples were used as templates for quantitative amplification (qPCR) of 16S rRNA genes and the functional genes of *narG, nirK, nirS* and *nosZ* which were performed on a Light Cycler 96 system (Roche, Basel, Switzerland) using primers Uni331F/Uni797 (Nadkarni et al., 2002), *narG*-f/r (Bru et al., 2007), *nirK*1040/FlaCu (Henry et al., 2004), *nirS*cd3A/R3cd58 (Kandeler et al., 2006) and *nosZ*-2f/2r61 (Henry et al., 2006) respectively. Detailed PCR reactions and amplification condition were performed as previously described (Yang et al., 2017).

### Community analysis by 16S rRNA gene sequencing and PICRUST

A sequencing library of V3-V4 regions of the 16S rRNA gene amplicons was established by two-step PCR amplification according to Illumina’s instructions as described in the previous literature (Wu et al., 2019) to investigate the bacterial community diversity and structure. Primer sets of 341F and 805R with Illumina MiSeq adaptor sequences were used in the first PCR amplification. After gel purification, the purified PCR amplicons were subjected to an additional round of amplification (index PCR) to label the individual samples by index sequence. Purified amplicons were sequenced on Illumina MiSeq System (Illumina Inc., United States). The quality controlled raw data were trimmed and analyzed as described previously (Wu et al., 2019). Representative operational taxonomy units (OTUs) were selected by UPARSE’s default (Edgar, 2013). In addition, reference-based chimera detection was performed using UCHIME (Edgar et al., 2011) against the RDP classifier training database (v9) (Cole et al., 2014). Finally, the OTU table was generated by mapping quality-filtered reads to the obtained OTUs with the Usearch (Edgar, 2010) global alignment algorithm at a 97% cutoff. Representative sequences for each OTU were submitted to the online RDP classifier (RDP database version 2.10) to determine the phylogeny with a bootstrap cutoff of 80%. The sequences of all the samples were downsized to 12000 to normalize the sequencing depth of all samples. Further analysis of alpha and beta diversity was performed using the QIIME platform (version 1.8) (Caporaso et al., 2010).

The functional gene compositions were predicted based on 16S rRNA gene information by using Phylogenetic Investigation of Communities by Reconstruction of Unobserved States (PICRUSt) (Langille et al., 2013). Tridimensional principal coordinates analysis (PCoA) based on the Bray-Curtis distance matrices and the clustering of the different groups was performed using MATLAB 2018a (The MathWorks Inc., Natick, MA, USA) to demonstrate the difference of total microbial community and denitrifying bacteria community in BF and FF. Variation significance among the different groups was conducted with multivariate analysis of variance (MANOVA) in MATLAB 2018a. Linear discriminant analysis (LDA) effect size (LEfSe) analysis was performed using parameters of p<0.05 and LDA score 3.5 (Segata et al., 2011). Spearman’s rank correlation test using MATLAB 2018a was performed to explore the correlation between the relative abundance of the *norB*- or *nosZ*-containing key OTUs and N_2_O/(N_2_O+N_2_). Spearman correlation analysis between soil properties and denitrifying functional genes were performed by using R. The statistical differences of all measured data in soil samples were analyzed with two-way ANOVA by Prism 6.0 (GraphPad, California, USA).

### Isolation of the most predominant denitrifying bacteria

Three grams of BF or FF from the N250+G treatment were diluted 1:10 in 8.5% NaCl and shaken for 30 min at 180 rpm and 28 °C. Five duplicate plates were spread individually by six gradient diluted soil suspensions from 10^−2^ to 10^−7^ dilution on 1/10 TSA medium (Tryptic Soy Agar, Merck, Darmstadt, Germany) and incubated under both aerobic and anaerobic conditions at 28 °C for two weeks, and inspected regularly to detect newly appearing colonies. Single, well-spaced colonies were selected based on morphology, aiming at obtaining as large a variation of bacterial isolates as possible. The picked colonies were further purified by streaking on new plates with the same medium.

### Identification of the denitrifying bacteria

The DNA samples of isolates were amplified to screen *Rhodanobacter*-like isolates by using universal eubacterial primers Rhodano-381F (5’-CAAGCCTGATCCAGCAATG-3’) and Xantho-817R (5’-CAACATCCAGTTCGCATCG-3’) (Green et al., 2012). The strain of *Castellaniella* enriched in FF was screened by high-throughput identification on partial sequencing of 16S rRNA gene (Zhang et al., 2019). Targeted bacterial isolates belonging to the predominant denitrifying bacteria were further taxonomically classified based on their full length 16S rRNA gene sequences. Each isolate was cultivated in 1/10 TSB medium. DNA was extracted using standard procedures (Omega E.Z.N.A. Bacterial DNA Kit). Enterobacterial repetitive intergenic consensus PCR (ERIC-PCR) was conducted for the genotyping of all isolates (Wei et al., 2004). Representative strains for different types of ERIC fingerprints were selected for 16S rRNA gene amplification using universal eubacterial primers 27F (5’-AGAGTTTGATCCTGGCTCAG-3’) and 1492R (5’-GGTTACCTTGTTACGACTT-3’) (Baker et al., 2003) and sequencing to identify the taxonomy. Each vial contains 30 ml of medium which was described in previous literature (Liu et al., 2013) with 0.505 g l^−1^ KNO_3_ (only for bacteria isolates of *Castellaniella*)or 0.1 g NaNO_2_ (only for bacteria isolates of *Rhodanobacter*) as an electron acceptor was inoculated with isolate inoculum and measured the denitrifying ability of bacterial isolates using Robot system.

### Inoculate *Castellaniella* sp. OFA38 into BF and FF

The *Castellaniella* sp. OFA38 liquid culture in TSB medium were centrifugally collected, suspended in Ringer solution (sodium chloride 9 g, potassium chloride 0.4 g, anhydrous calcium chloride 0.25g, dissolved in 1L pure water), and added to BF and FF with 10^7^ cells/g soil. The headspace of serum vials was sampled every 4 hours and analyzed for N_2_ and N_2_O concentration in a robotized incubation system similar to that described by Molstad (Molstad et al., 2007) until no more N_2_ produced.

### Accession numbers of the sequence data

The raw Illumina sequence data generated in this study have been deposited to the GenBank Sequence Read Archive (SRA) database in the National Center for Biotechnology Information (NCBI) under the accession number PRJNA755188. 16S rRNA gene sequence for selected predominant bacterial strains were deposited in GenBank under accession numbers MZ824722-MZ824747.

## Results

### Physicochemical properties of soil samples

The soils of BF and FF had different physicochemical properties, as shown by the indices (Table 1). The water content, WHC, DOC and ammonium level of BF were higher than those of FF, but the pH, PDR and nitrate values of BF were lower than those of FF. Most nitrate remained in the samples of the N250 group at the end of anaerobic incubation (Table S1), and this group showed a decrease in pH. In contrast, supplementation with glucose resulted in an approximately complete reduction in nitrate and increased pH in both soils. There was no accumulation of nitrite in any incubation setting except for a low level in the N250 group of BF. In addition, the ammonium level in BF was higher than that in FF regardless of conditions.

### Gas kinetics during anaerobic incubation

The N_2_O concentrations in the vials of CK for both BF and FF were very low throughout the incubation period compared to those of the N250 and N250+G groups. The N_2_O kinetics in BF and FF showed a similar trend in CK, with the N_2_O concentration slowly peaking and then decreasing during the incubation period. No obvious NO_2_ accumulation in FF was observed in CK after 64 hours (Fig. 1a). In the N250 group, the N_2_O concentration in BF constantly increased during the incubation period, while it slightly increased early on but gradually declined after 96 h of incubation in FF (Fig. 1b). In the N250+G group, N_2_O accumulation in BF increased drastically before 80 h and thereafter quickly decreased. However, much less N_2_O accumulated in FF, and levels gradually decreased after 112 hours (Fig. 1c).

**Fig. 1.**
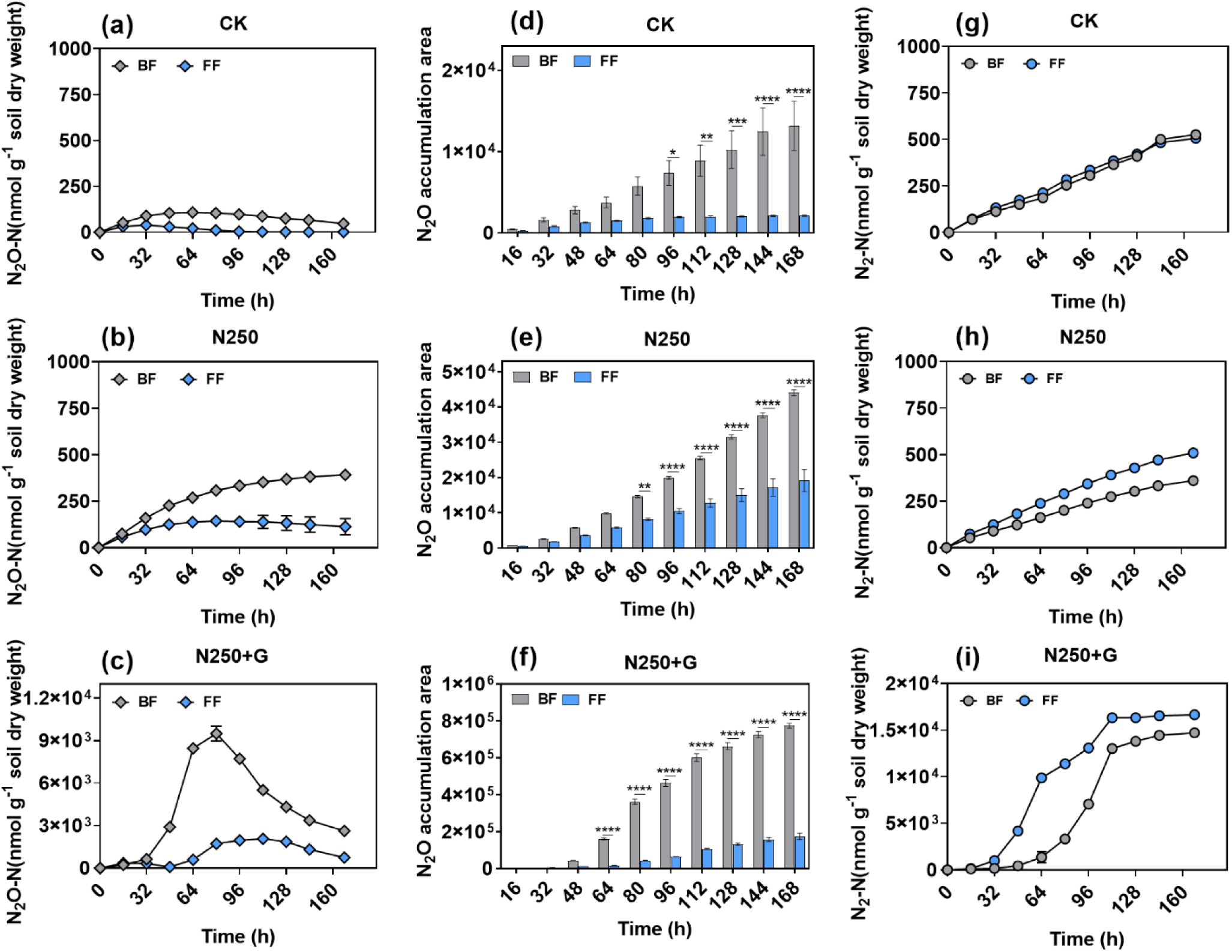
Kinetics of N_2_O and N_2_ during anaerobic incubation. Kinetics of N_2_O (**a-c**), N_2_O accumulation area (**d-f**) and kinetics of N_2_ (**g-i**) under three different treatments. Bars indicate means, and error bars indicate the SEM. Differences in the N_2_O accumulation area were calculated between BF and FF via a two-way ANOVA. *P<0.05, **P<0.01, ****P<0.0001.

The difference in the accumulation area under the N_2_O dynamic curve in BF and FF was significant (P<0.037, two-way ANOVA) after 96 h of incubation in all treatments. The final N_2_O accumulation area in BF with three different treatments was 2.3-6.2 times higher than that in FF (Fig. 1d-f).

The accumulation N_2_ production by BF and FF in the CK and N250 groups increased linearly during the incubation period (Fig. 1g-h). In CK, there was no difference in N_2_ accumulation between these two types of soil, while in the N250 and N250+G groups, N_2_ accumulation in FF was higher than that in BF. A large amount of N_2_ was released in both soils under sufficient carbon and nitrogen, although FF was consistently higher than BF over 168 h of incubation (Fig. 1i).

In addition, the area under the dynamic curve of N_2_O+N_2_ produced in BF and FF increased over time throughout the incubation period (Fig. S1a-c). The N_2_O/(N_2_O+N_2_) ratio of each type of soil was significantly stimulated by nitrate addition, whereas it declined with nitrate and glucose addition (Fig. S1d). Nitrate and nitrite levels at the end of a 7-day anaerobic incubation period were positively correlated with N_2_O/(N_2_O+N_2_) (Fig. S3a and b).

In general, N_2_O accumulation and N_2_O/(N_2_O+N_2_) in BF were consistently higher than those in FF regardless of the initial levels s of nitrate and glucose.

### Quantity of denitrifying and 16S rRNA genes

Quantitative polymerase chain reactions (qPCR) targeting the 16S rRNA gene and denitrifying genes revealed these genetic variations among the samples of different treatments. The copy numbers of the 16S rRNA (Fig. 2a), *nirK* (Fig. 2c) and *nosZ* (Fig. 2d) genes were significantly higher in FF than in BF in the CK and N250+G groups. These three genes were also higher in FF in the N250 group, although the difference was not significant (P>0.05). The copy numbers of the *narG* and *nirS* genes (Fig. 2b and c) were significantly higher in FF than in BF for all treatments. For the samples of the CK and N250 groups, the ratio of *narG*/*nosZ* (Fig. 2e) was not significantly different in BF and FF. However, in the N250+G group, the ratio was significantly higher in BF than in FF. The summed values of *nirK*/*nosZ* and *nirS*/*nosZ* (Fig. 2f) showed no significant differences in BF and FF in all treatments.

**Fig. 2.**
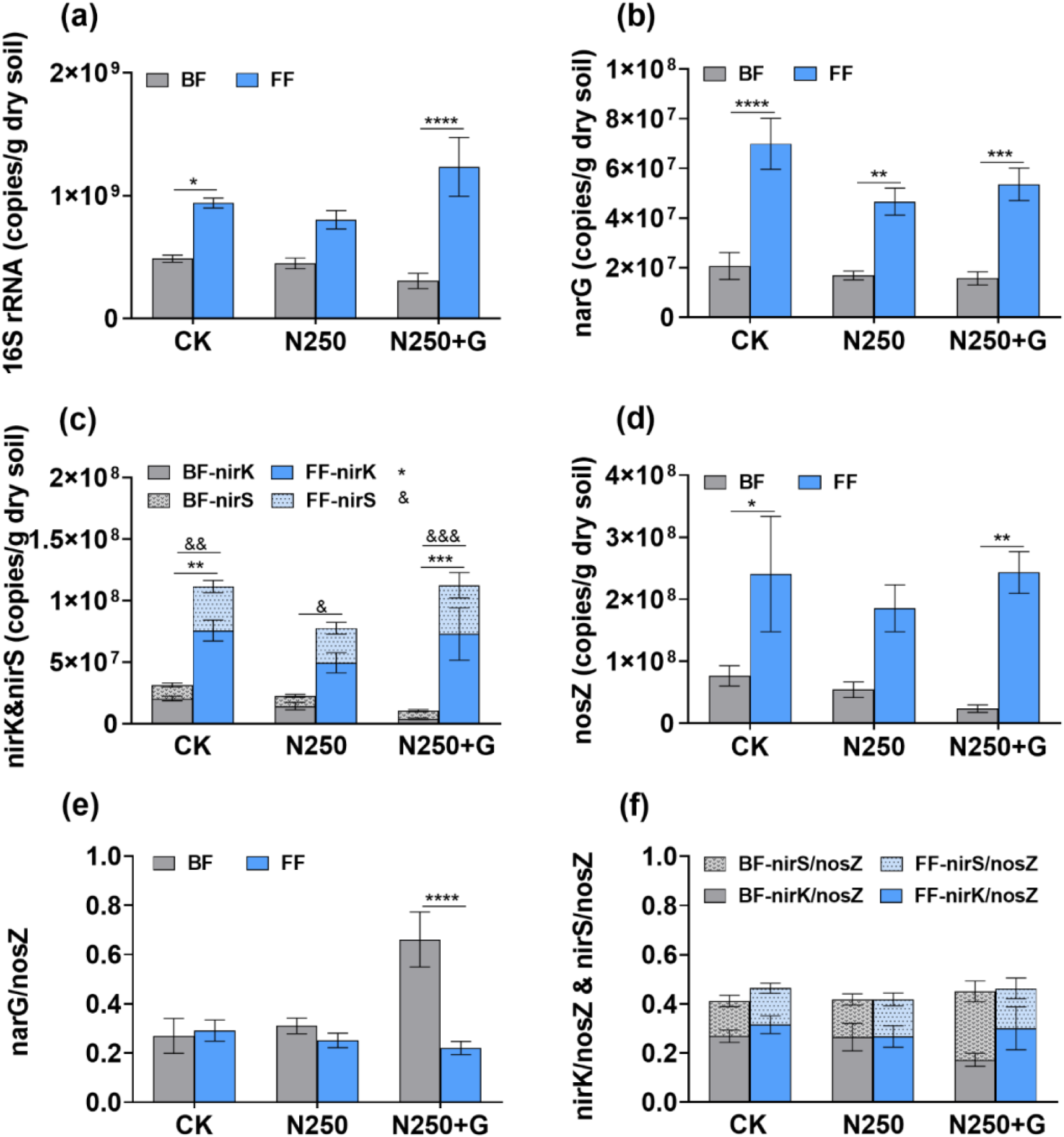
Copy number of genes involved in nitrogen cycling and ratio of denitrifying functional genes to *nosZ*. 16S rRNA (**a**), *narG* (**b**), *nirK* and *nirS* (**c**), *nosZ* (**d**). *narG*/*nosZ* (**e**), and *nirK*/*nosZ* and *nirS*/*nosZ* (**f**) of BF and FF with three different treatments. Bars indicate means, and error bars indicate the SEM. Differences in three kinds of ratios were calculated via a two-way ANOVA. *P<0.05, **P<0.01, ****P<0.0001.

Moreover, the ratios of *narG/*16S rRNA (Fig. S2a), *nirK/*16S rRNA (Fig. S2b), *nirS/*16S rRNA (Fig. S2c) *and nosZ/*16S rRNA (Fig. S2d) in all treatments showed no significant difference between BF and FF, except that the ratios of *nirK/*16S rRNA in FF of the CK and N250+G groups were significantly higher than those in BF. The N_2_O/(N_2_O+N_2_) ratio was negatively correlated with *nirK*/*nosZ* and *nosZ* but positively correlated with *nirS*/*nosZ* in both BF and FF (Fig. S3a and b). However, N_2_O/(N_2_O+N_2_) and *narG*/*nosZ* were positively correlated in BF but negatively correlated in FF.

### Variations in microbial community structures

In 18 samples, a total of 364,452 high quality 16S rRNA gene sequences were clustered into 6537 representative OTUs. There was no significant difference in alpha diversity (the Shannon index and Chao 1 index) between BF and FF, except for Chao 1 in the N250 group of FF, which was significantly higher than that of BF (Fig. S4a and b). The trajectory of the microbial community structure in the three-dimensional PCoA plot based on the Bray-Curtis distance showed divergence between the bacterial communities of BF and FF and the influence of carbon and nitrogen addition on the bacterial communities (Fig. 3a). MANOVA results confirm that the structures of the microbial communities of the two soils were significantly separated (****P<0.0001, MANOVA test). The microbial community structures of the CK and N250 groups were more similar (Fig. 3b) but significantly different from those of the N250+G group (***P<0.001, ****P<0.0001, MANOVA test).

**Fig. 3.**
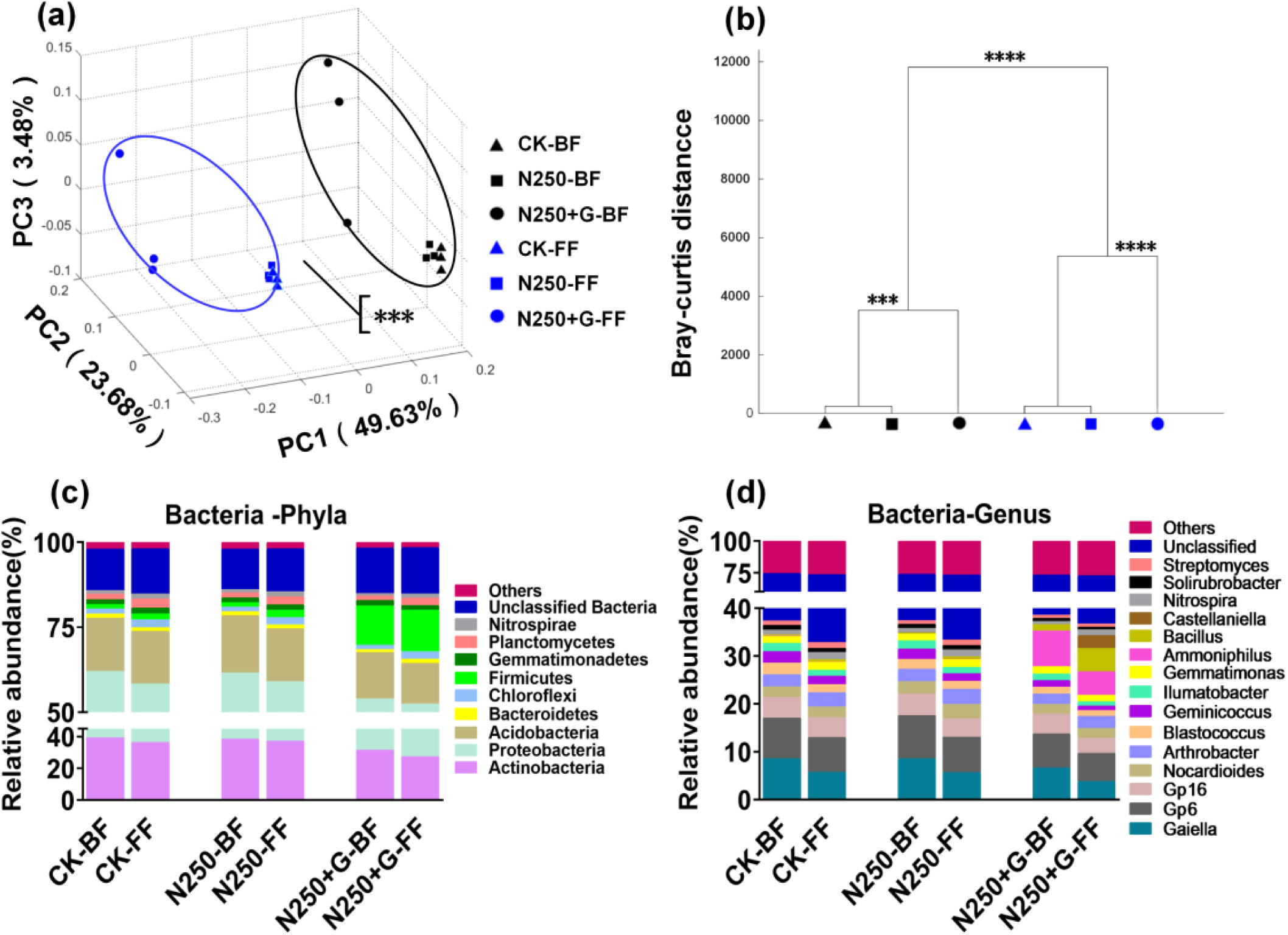
Alteration of microbial community structure and composition in BF and FF under three different treatments. Tridimensional PCoA plot of the microbial community based on the Bray-Curtis distance (**a**), ***P<0.001. Clustering of the microbial community based on Bray-Curtis distance calculated with a MANOVA test (**b**). ***P<0.001, ****P<0.0001. Distribution of dominant taxa of the microbial community in BF and FF at the phylum level (**c**) and at the genus level (**d**). The predominant phyla or genera with a relative abundance higher than 1% are displayed separately.

There were nine dominant phyla, including Actinobacteria, Proteobacteria, Acidobacteria, Bacteroidetes, Chloroflexi, Firmicutes, Gemmatimonadetes, Planctomycetes and Nitrospirae, except for 12-13.6% of all OTUs, which were unclassified bacteria (Fig. 3c). Notably, glucose specifically enriched bacteria belonging to Firmicutes.

The dominant genera (relative abundance ≥1%) in BF and FF were measured at approximately 37.3~38.6% and 32.9~36.8%, respectively, including *Gaiella, Gp6, Gp16, Nocardioides, Arthrobacter, Blastococcus, Geminicoccus, Ilumatobacter, Gemmatimonas, Ammoniphilus, Bacillus, Castellaniella, Nitrospira, Soilrubrobacter* and *Streptomyces* (Fig. 3d). The relative abundance of these genera in each type of soil was influenced by carbon and nitrogen sources. In particular, when supplemented with nitrogen and glucose, *Ammoniphilus* was enriched in both BF and FF, while *Castellaniella* and *Bacillus* were more enriched in FF than in BF.

### Predicting denitrification function and functional bacteria of soil communities

In all samples, 1197 and 732 of 6537 OTUs were predicted for the *norB* and *nosZ* genes, respectively. The Shannon index for *norB-*containing denitrifying bacteria in BF was significantly higher than that in FF in the presence of sufficient carbon and nitrogen, although the values in the two types of farmland soils in the CK and N250 groups were not significantly different (Fig. S4c). It is worth noting that the Shannon diversity of *nosZ* in FF was significantly higher than that in BF under all treatments (Fig. S4e). However, the Chao 1 index for *norB-* (Fig. S4d) or *nosZ-* (Fig. S4f) containing denitrifying bacteria in both soils exhibited no difference (two-way ANOVA).

The Bray-Curtis distance showed that the community structures of *norB*- (Fig. S5a and b) or *nosZ*- (Fig. S5e and f) containing bacteria in BF and FF were significantly different, while the beta diversity within the same soil showed that the CK and N250 groups were significantly different from the N250+G group (P<0.0001, MANOVA test).

### *norB*-containing bacteria in two types of soils

A total of 1197 *nor*B-containing OTUs were affiliated with 3 dominant phyla, namely, Acidobacteria, Actinobacteria and Proteobacteria, as well as other rarer phyla (relative abundance of phylum <1%) (Fig. S5c).

At the genus level, there were 12 dominant *norB*-containing genera (Fig. S5d). Among these genera, 7 lacked the *nosZ* gene. Five of these 7 genera, *Candidatus Solibacter, Dokdonella, Kaistobacter, Lysobacter* and *Phyllobacterium* were more abundant in BF than in FF. Another two genera, *Thermomonas* and *Castellaniella*, were more abundant in FF than in BF in the N250+G group. *Azospirillum, Dechloromonas, Devosia* and *Rhodonobacter* among the predominant bacterial genera predicted to contain both *norB* and *nosZ* were more abundant in BF than in FF. In contrast, genus *Azoarcus* containing both *norB* and *nosZ* was more enriched in FF than in BF when sufficient carbon and nitrogen sources were added. However, it should be noted that more than 60% of *norB* denitrifiers were not affiliated with any known genus.

At the OTU level, 39 key OTUs selected by LEfSe could be used as indicators to distinguish the *norB* denitrifying communities between BF and FF (Fig. S6a-c). Among them, only OTU114930 of *Azospirillum*, which was enriched in BF in N250+G, showed a negative correlation with the N_2_O/(N_2_O+N_2_) ratio, while the other 21 OTUs enriched in BF (especially the OTUs affiliated with *Kaistobacter* and *Rhodonobacter* as predominant bacteria in BF) showed a positive correlation with the N_2_O/(N_2_O+N_2_) ratio (Fig. 4a). In contrast, 17 OTUs enriched in FF showed a negative correlation with the N_2_O/(N_2_O+N_2_) ratio.

**Fig. 4.**
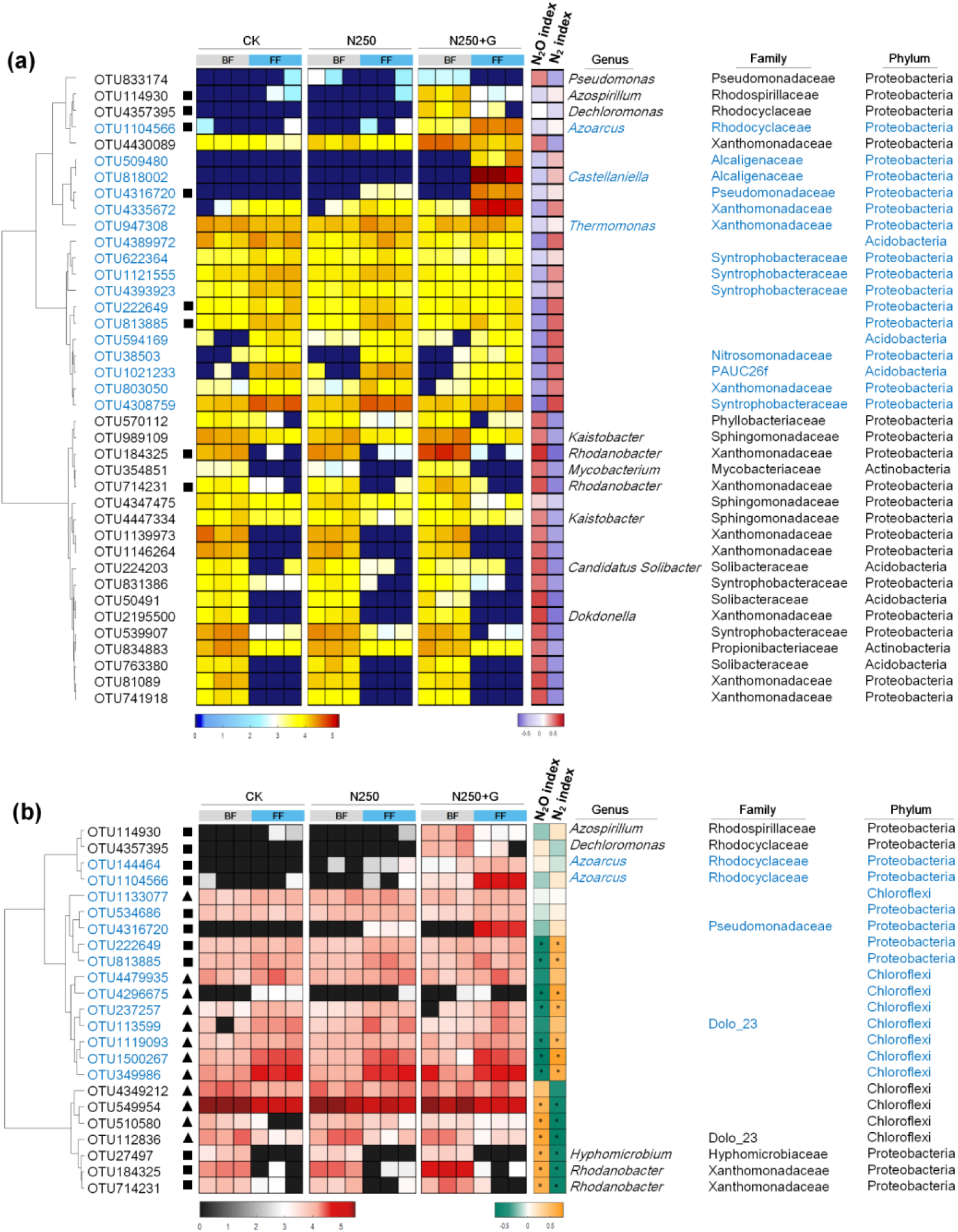
Heatmap of key denitrifiers identified in BF and FF with different treatments. Heatmap of the 39 key OTUs containing *norB* that differed between the two types of soils (**a**). Heatmap of the 23 key OTUs containing *nosZ* showing differences between the two types of soils (**b**). OTUs marked with ‘■’ represent those containing both *norB* and *nosZ*, whereas OTUs marked with ‘▲’ represent OTUs containing *nosZ* but lacking *norB*. OTUs without marks represent OTUs containing *norB* but lacking *nosZ*. The colours of the spots in the left panel represent the relative abundance (normalized and log-transformed) of the OTUs in each sample, while in the right panel denote the R-value of the Spearman’s correlation between the OTU and N_2_O/(N_2_O+N_2_) and N_2_/(N_2_O+N_2_).

### *nosZ-*containing bacteria in two types of soils

A total of 732 *nosZ*-containing OTUs were affiliated with Bacteroidetes, Chloroflexi, Proteobacteria, Verrucomicrobia and other phyla (relative abundance of phylum <1%) (Fig. S5g). The relative abundance of 7 dominant genera, *Rhodanobacter, Opitutus, Devosia, Dechloromononas, Caldilinea, Azospirillum* and *Azoarcus* varied in BF and FF under the different treatments (Fig. S5h). *Caldilinea* and *Opitutus* containing *nosZ* but lacking *norB and Azoarcus* containing both *nosZ* and *norB* were more abundant in FF than in BF when sufficient carbon and nitrogen were provided. More than 81% of *nosZ* denitrifiers could not be affiliated at the genus level.

At the OTU level, 23 OTUs were identified as key members that contributed to the discrepancy in the *nosZ* denitrifying community found between BF and FF (Fig. S6d and f). Two OTUs of *Rhodanobacter*, an OTU of *Hyphomicrobium*, and three OTUs of Chloroflexi were enriched in BF and significantly positively correlated with the N_2_O/(N_2_O+N_2_) ratio, while several OTUs of Chloroflexi and Proteobacteria were enriched in FF and significantly negatively correlated with the N_2_O/(N_2_O+N_2_) ratio (Fig. 4b).

### Denitrification functions of dominant bacteria in BF and FF

Among the 437 isolates from BF, 25 were identified as bacteria belonging to genus *Rhodonabacter*. These isolates were clustered into 18 strains by ERIC typing. Fifteen of these strains had 100% 16S rRNA gene sequence similarity with the most predominant *Rhodonabacter* OTU in the BF bacterial community, while the remaining 3 strains had high sequence similarity with another much less abundant *Rhodonabacter* OTU (Table S2). Seventy-two of the 456 isolates obtained from FF belonged to *Castellaniella* and were clustered into 8 different strains by ERIC typing. All of these strains had 100% 16S rRNA gene sequence similarity with the *Castellaniella* OTU in the FF (Table S3).

The denitrifying functions of representative strains of these two genera were measured. Eighteen representative isolates of *Rhodanobacter* were only capable of reducing nitrite, rather than nitrate, to produce N_2_O without any further reduction in most strains corresponding to the most abundant OTU in the BF or only slight reduction in a few strains corresponding to the less abundant OTU, resulting in the obvious accumulation of N_2_O (Fig. 5a). However, all representative isolates of *Castellaniella* effectively reduced nitrate to N_2_ without any accumulation of N_2_O (Fig. 5b).

**Fig. 5.**
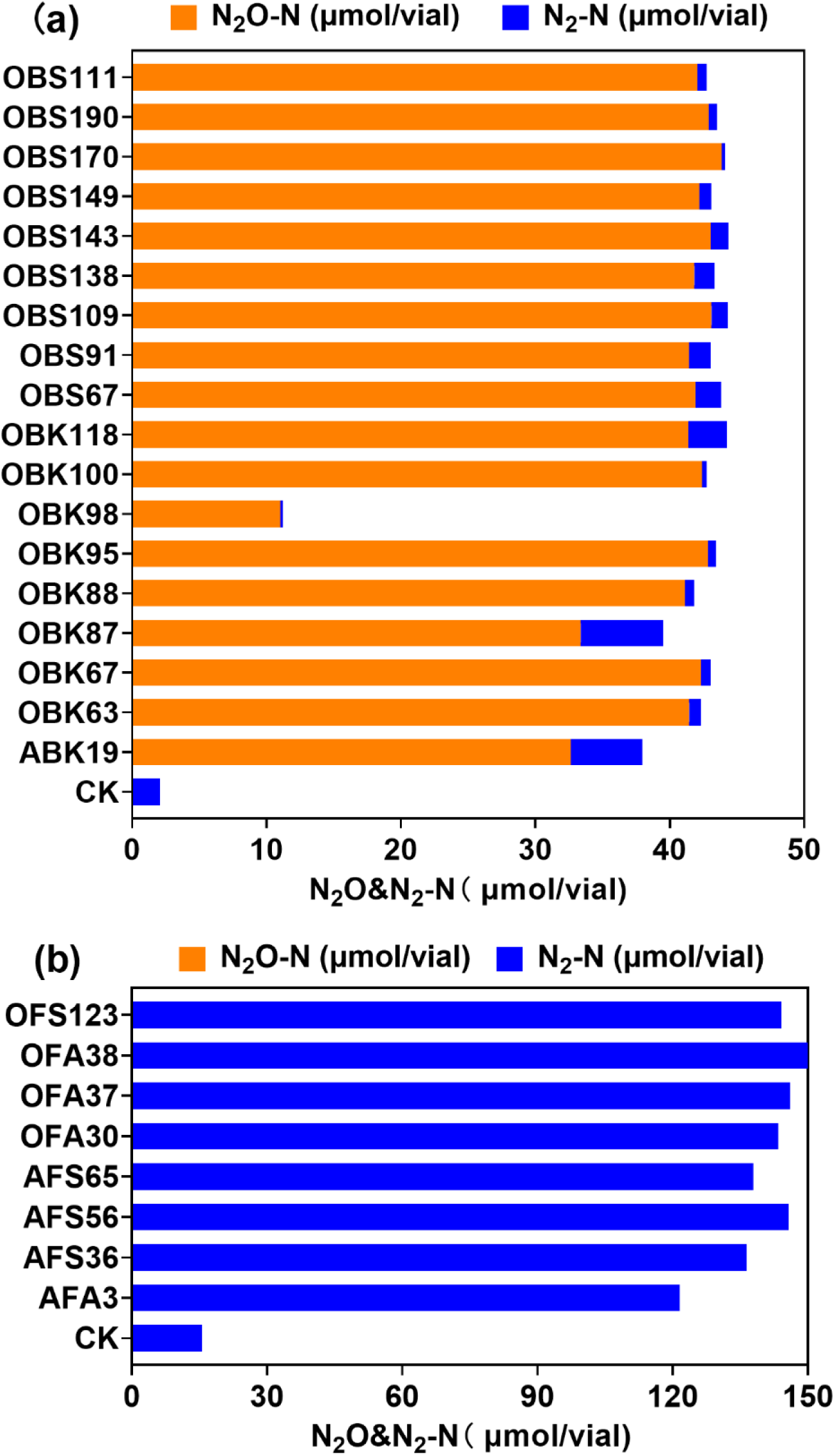
Denitrification functions of 18 strains of *Rhodanobacter* isolated from BF in DM medium containing 43.5 μmol/vial nitrite (**a**) and 8 strains of *Castellaniella* isolated from FF in DM medium containing 149.9 μmol/vial nitrate (**b**).

### N_2_O reduction of BF and FF inoculated with the isolate of *Castellaniella*

*Castellaniella* sp. OFA38 is one of the 8 different strains of *Castellaniella* isolated from FF, which rapidly reduced N_2_O to N_2_ and efficiently reduced N_2_O emissions in the microcosm experiment performed using BF (Fig. 6a). During the incubation period of 64 h, the N_2_O accumulation area in inoculated BF decreased by 97.5% compared to that in the same soil without inoculation. Since very little N_2_O accumulated in the uninoculated FF, inoculation with *Castellaniella* sp. OFA38 in this soil showed only a marginal effect on N_2_O accumulation, which was 26.4% lower than that observed in uninoculated FF at 64 h (Fig. 6b).

**Fig. 6.**
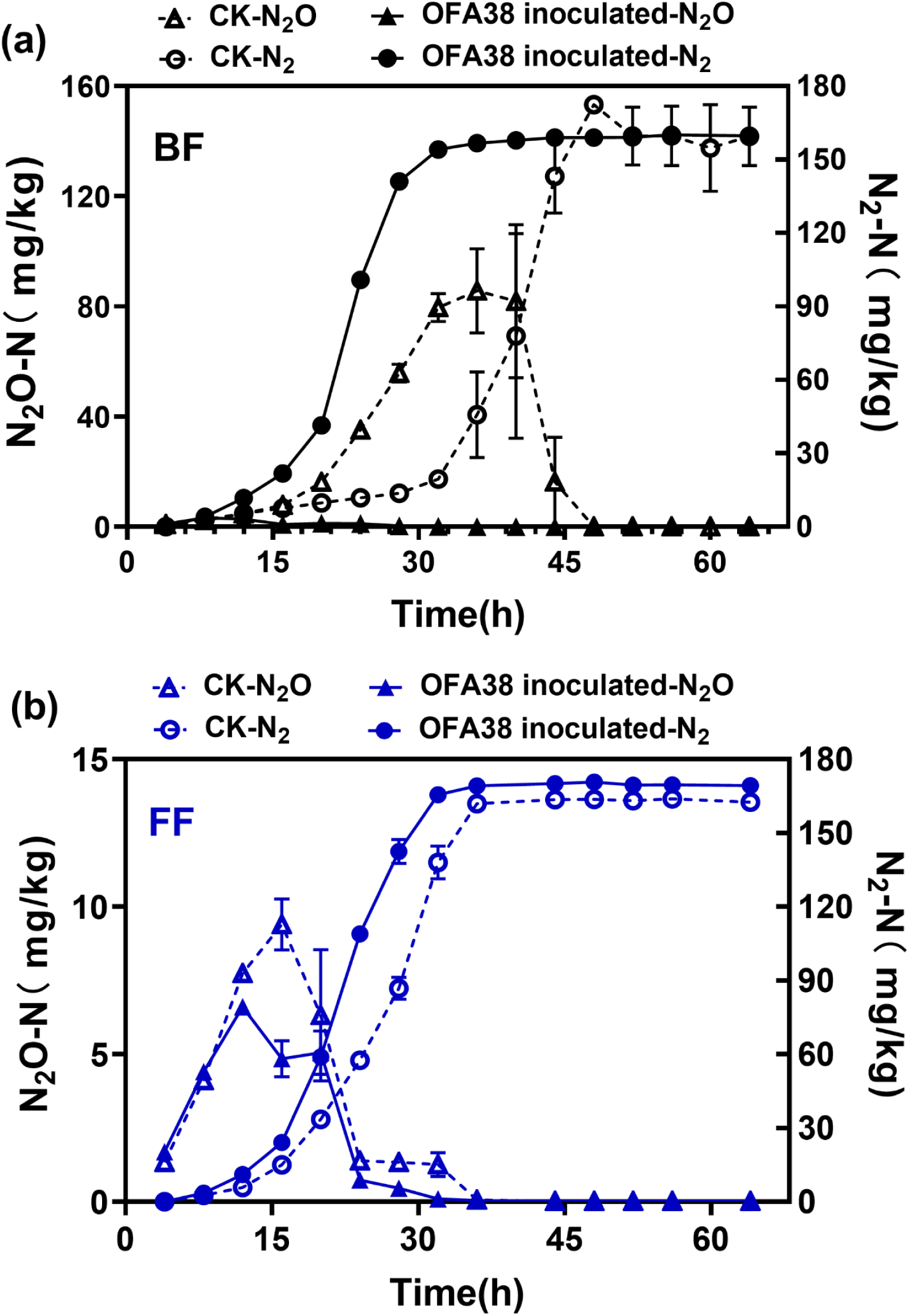
Effects of *Castellaniella* sp. OFA38 on N_2_O emissions from BF (**a**) and FF (**b**) during anaerobic incubation.

## Discussion

### Effect of nitrogen and carbon on N_2_O accumulation

The nitrate in BF and FF was almost completely reduced to N_2_O and N_2_ over 7 days of incubation in the N250+G group, which is consistent with a pulse emission of N_2_O produced due to strong denitrification immediately after fertilization and irrigation in the field (Whalen, 2000) and which shows that both C and N might be limiting factors of denitrification activity in these two types of soils (Wu et al., 2013). N_2_O accumulation had already peaked before 7 days in all treatments. In addition, we intended to use the activity potential of two soils with their original microbial community without any shift of structures to represent the in situ microbial activity. Therefore, we incubated and monitored the nitrogen gases for only 7 days. Measuring denitrification under different carbon and nitrogen conditions would help fully elucidate the relationship between the soil microbiome and N_2_O emissions.

Many studies indicate the effects of nitrogen and carbon on denitrification. In terms of each type of soil, there was no difference in the microbial community composition of the CK and N250 groups, but the amount of N_2_O produced in the N250 group was higher than that in the CK group because pulses of NO_3_^−^ may have a substantial and immediate impact on N_2_O emission without a prior change in denitrifying community structure (Carrino-Kyker et al., 2012). A few previous studies show that the reducing power for nitrogen reduction is generated from carbon oxidative catabolism from the perspective of individual metabolism (Zumft, 1997; Dias et al., 2009). Therefore, the accumulation of N_2_O during denitrification is a result of an imbalance in carbon and nitrogen metabolism (Dai et al., 2020). As expected in this study, much more N_2_O accumulated in the group with added glucose. Glucose significantly changed the results of the microbial community, promoted high N_2_O emissions, and then gradually reduced N_2_O in each type of soil, because glucose, as a simple C substrate, can be easily utilized by microorganisms and increases soil microbial activity, leading to NO_3_^−^ depletion and an increased consumption of N_2_O (Miller et al., 2008).

Nevertheless, nitrate and DOC levels were not direct proximal factors determining the difference in N_2_O accumulation between BF and FF. Relative to BF, FF showed a higher efficiency in consuming N_2_O. In addition, the difference in N_2_O accumulations in BF and FF was more obvious when higher levels of nitrogen and carbon were presented.

### Effect of denitrifying gene abundance on N_2_O accumulation

Previous studies have demonstrated that the N_2_O flux is correlated with the quantity of denitrifying genes (Wang et al., 2013; Li et al., 2016; Li et al., 2017). For example, one study found that the annual N_2_O emissions in intensively managed calcareous FF to have a significant correlation with the abundance of denitrifying genes, such as the *narG, nirK* and *nirS* genes (Yang et al., 2017). In addition, another study suggests that a strong correlation exists between *nirS* gene abundance and potential N_2_O emissions (Cui et al., 2016). However, the accumulation of N_2_O is not simply determined by how much is produced but is also related to the reduction of N_2_O. Higher rates of both the production and reduction of N_2_O do not result in more accumulated N_2_O because N_2_O emissions are the result of a net balance between their production and consumption (Maeda et al., 2011). Recently, numerous studies have highlighted that substantially higher N_2_O emissions are attributed to the relative abundance of denitrifying bacteria either generating (NO_3_^−^→N_2_O, primarily catalysed by nitrite reductase encoded by *nirS*/*nirK*) or degrading N_2_O (N_2_O → N_2_, catalysed by N_2_O reductase encoded by *nosZ*) (Regan et al., 2011), such as the activity ratios of NIR/NOS and NOR/NOS (Ye et al., 2021).

Our findings indicate that the quantity of denitrifying functional genes *narG, nirK, nirS* and *nosZ* was generally higher in FF, which implies that potential for the production and reduction of N_2_O in FF was stronger than that in BF. Nevertheless, the quantity of denitrifying genes and lack of differences between the ratios of denitrifying genes of the two types of soils do not explain the consistently higher levels of accumulated N_2_O in BF than in FF. Similar phenomena have been reported by several other studies, such as studies on dazomet-fumigated FF and lateritic red soil (Fang et al., 2018) and on differently fertilized soils in a vegetation greenhouse (Ji et al., 2021).

### Effect of denitrifier composition on N_2_O accumulation

The denitrifying microbial communities in agricultural soils are very diverse and complex in nature. Numerous studies have revealed that N_2_O emissions are anchored in the taxonomic composition of denitrifier communities (Braker et al., 2012; Cui et al., 2016; Ji et al., 2021). For example, one study suggests that the taxonomic makeup of bacterial communities, but not the size of the communities determines the level of denitrification activity (Braker et al., 2015). In addition, Liu et al. founded eight *Thauera* strains of denitrifying bacteria to be divided into two distinct denitrification regulatory phenotypes by comparing the denitrification dynamics of these strains based on the accumulation of NO_2_^−^, NO and N_2_O, oxic/anoxic growth and transcription of functional genes (Liu et al., 2013). This result shows that microorganisms from the same genus might have different denitrification functions. A previous study found that different species of denitrifying bacteria in samples may play an important role in regulating denitrification (Tilman et al., 2014). Therefore, the structures of the denitrifying community are of great importance to the feedback of potential N_2_O emissions (Rich et al., 2003).

The results of this study clearly indicate that the different denitrifying communities in the two studied soils played a crucial role in regulating N_2_O. First, we found a higher diversity of *nosZ* but a lower diversity of *norB* in FF than in BF, as showed by the Shannon index. This may be beneficial for N_2_O reduction in FF and N_2_O production in BF due to the presence of more diverse denitrifiers, including some highly efficient N_2_O-reducing bacteria in FF. Previous studies have also showed that microbial diversity plays a role in N_2_O emissions (Philippot et al., 2013; Wagg et al., 2014). Second, N_2_O-generating bacteria containing *norB* but lacking *nosZ* were found at much higher levels in BF than in FF, implying the presence of more net N_2_O-producing bacteria in BF. In this study, the abundance of the dominant bacteria of *Kaistobacter* that produce N_2_O but lack the *nosZ* gene for N_2_O reduction was invariably higher in BF than in FF under different carbon and nitrogen sources. In particular, the two OTUs of *Kaistobacter* enriched in BF contributed to the structural divergence of the *norB*-containing bacterial community in BF and FF and were positively correlated with N_2_O/(N_2_O+N_2_). This may be a reason for the high levels of accumulated N_2_O found in BF, and the same result has been reported for farmland soil with conventional intensive N fertilization (Ji et al., 2021). Similarly, specific N_2_O-reducing bacteria were dominant in FF. Therefore, these denitrifiers in the soil determined the efficiencies for the production of N_2_O and reduction of N_2_O to N_2_. This conclusion is also consistent with a previous study indicating that increased N_2_O emissions in Ekhaga soils are accompanied by an increase in the amount of inoculated denitrifying bacteria *Agrobacterium tumefaciens* C58, which lacks the *nosZ* gene, establishing a direct causal link between the composition of the denitrifier community and potential N_2_O emissions (Philippot et al., 2011). Furthermore, based on the analysis of the 16S rRNA gene sequences, we identified several key denitrifiers. For each type of farmland soil, glucose significantly altered the composition of soil denitrifying bacteria, whereas strains of *Rhodanobacter* as predominant bacteria in BF were constantly present under different carbon and nitrogen conditions and positively correlated with N_2_O/(N_2_O+N_2_). Similar phenomena have been reported for conventionally fertilized soil (Ji et al., 2021). However, *Castellaniella* was enriched in FF under sufficient carbon and nitrogen conditions. Numerous studies have shown that microorganisms of *Rhodanobacter* and *Castellaniella* play important roles in denitrification.

A previous study founded *Rhodanobacter* to be the most abundant genus among all denitrifiers in the soil samples studied on a chronic fertilization experiment performed in a black agricultural soil field (Hu et al., 2020). The unique physicochemical properties of fertilized BF probably establish a niche for highly abundant *Rhodanobacter*, as was observed for BF in this study. Bacteria from *Rhodanobacter* were identified as acid-tolerant denitrifiers, and their distribution was driven by pH (Green et al., 2012). Studies have shown that strains of *Rhodanobacter* isolated from terrestrial subsurface sediments and from nitrate-rich zones of a contaminated aquifer are complete denitrifiers (Green et al., 2010; Prakash et al., 2012), but N_2_O significantly accumulates during denitrification (Green et al., 2010). In another study, the *Rhodanobacter*-like bacterial community demonstrated significant denitrification with N_2_O as an end-product at a low pH and under controlled conditions (Van den Heuvel et al., 2010). In addition, three truncated denitrifiers of *Rhodanobacter* isolated from the acidic soil of Fjaler on the western coast of Norway, accumulated N_2_O under denitrifying conditions, although one was capable of N_2_O reduction (Lycus et al., 2017). Although the literatures report that bacteria from *Rhodanobacter* tend to accumulate N_2_O during denitrification, different strains of one genus may show different capacities for producing and consuming N_2_O (Nadeem et al., 2013; Liu et al., 2016). Therefore, the real denitrifying metabolism of *Rhodanobacter* which predominates in BF, still needs to be verified.

Bacteria from *Castellaniella* were also identified as key denitrifying bacteria in this study. Denitrification by strains of *Castellaniella* has been widely reported (Pang and Liu, 2007; Spain and Krumholz, 2011; Deng et al., 2021). For example, the *Castellaniella* bacterium effectively reduces nitrate to gaseous nitrogen, thus reducing the nitrate concentration in sewage (Pang and Liu, 2007). However, the N_2_O production and reduction capacities of *Castellaniella* in the soil environment have not been reported, especially for fertilized agricultural soil. In this study, we report the isolation of a denitrifying bacterium of *Castellaniella* that can rapid reduction of N_2_O without the accumulation of N_2_O, representing—the first observation of highly efficient N_2_O-reducing bacteria of *Castellaniella* in farmland soil.

### N_2_O metabolism of isolates corresponding to the key bacteria in soil

In this study, we found the PICRUSt prediction results to be inconsistent with our expectations. For example, *Castellaniella* was predicted to contain only the *norB* gene, and *Rhodanobacter* was predicted to contain both the *norB* and *nosZ* genes. However, these results do not explain the correlations of these bacteria with N_2_O levels in the two soils. Therefore, we isolated bacteria from the two soils by culturomics and screened denitrifying isolates belonging to these predominant genera for physiological function measurements. Among 18 *Rhodanobacter* isolates, only a few isolates corresponding to the less abundant OTU of *Rhodanobacter* slightly reduced N_2_O, and the remaining isolates could not reduce N_2_O. These bacteria could be the source of strong N_2_O accumulation in BF due to their high abundance. Eight isolates of *Castellaniella* obtained from FF contained highly active N_2_O reductase and did not accumulate N_2_O when incubated in nitrate-containing medium. These results indirectly prove that the predominant denitrifying bacteria of *Castellaniella* contributed to the efficient reduction of N_2_O in FF. In addition, the isolation of highly efficient N_2_O-reducing bacteria, such as the isolates of *Castellaniella* sp. OFA38 described in this study, rapidly reduced N_2_O to N_2_ and efficiently reduced N_2_O emissions in BF, exhibiting great potential to mitigate N_2_O emissions from farmland soil, even in FF where *Castellaniella* is already found in high abundance.

## Conclusions

This study demonstrates that two types of important Chinese agricultural soils show different N_2_O accumulation patterns during anaerobic incubation, with BF accumulating more N_2_O. Investigations of the microbial mechanism show that the difference in N_2_O accumulation is not due to the differences in nitrate and DOC contents or the quantity of denitrifying genes in soils. The different compositions of denitrifiers in the two types of soil could be the main reason for the difference in N_2_O accumulation. Most of the key *norB-*containing OTUs enriched in BF were found to be positively correlated with N_2_O/(N_2_O+N_2_), while other *norB*-containing and *nosZ*-containing key OTUs enriched in FF were found to be negatively correlated with N_2_O/(N_2_O+N_2_). As proven by isolation, bacteria of the most abundant members of *Rhodanobacter* in BF were only capable of reducing nitrite to N_2_O and accumulating N_2_O. Bacterial isolates of *Castellaniella* enriched in FF showed a capacity for the rapid reduction of N_2_O to N_2_. The results of this study provide a better understanding of the bacterial N_2_O source in BF and the bacterial N_2_O sink in FF. The findings of this study provide new insights into the strategies for mitigating N_2_O emissions in different agricultural soils, by regulating the composition of the denitrifier community.

## Acknowledgments

The authors would like to express their gratitude to Dr. Wei Zhang (Institute of Applied Ecology, CAS) for her help in collecting the black soil.

